# Mating signal energy gradient mediates gene flow in the early stage of avian speciation

**DOI:** 10.1101/2025.03.28.646037

**Authors:** Derek Daniel Eddo, Zachary James Hodur, Silu Wang

## Abstract

The grey zones of speciation harbor hybridizing species that experience the tension of incompatibility and gene flow. Mating signals for mate choice or competition in the hybridizing species can mediate heterospecific mating, thus predicting the extent and direction of gene flow in the entangled evolutionary trajectories. Some mating signals are diffusive, facilitating gene flow across species boundaries; some asymmetrically mediate gene flow from one species to another; while others prevent hybridization and strengthen species boundaries. Here we investigate the role of mating signals in mediating gene flow at nascent species boundaries. By comparatively studying the mating signals of 109 hybridizing avian species pairs, we discovered: (1) the visual, acoustic, kinesthetic, and chemosensory signaling modalities are associated with their roles in the species boundaries; (2) the relative energetic expenditure of mating signals predicts the directionality of gene flow between species; (3) the effects of mating signals on species boundaries are associated with mitochondrial genetic distance. This synthesis highlights the significant role of mating signal energetics in shaping the grey zone of avian speciation.

## Introduction

In the avian speciation continuum, mating signals often mediate hybridization as they stimulate behavioral changes of the receiver to increase the chance of reproduction of the signalers, even in heterospecific contexts (Endler 1992; Ryan 2018; de Zwaan et al. 2022). Mating signals penetrate various sensory dimensions including visual (Jones and Hunter 1998; Coster et al. 2018; Zonana 2019; Linck et al. 2020), acoustic (Jouventin et al. 2006; Marcondes et al. 2013; Augustine and Trauba 2015), kinesthetic (Hunter 1987; Fefelov 2001; Myers et al. 2019), and chemosensory modalities (Soini et al. 2006; Van Huynh and Rice 2019) (Fig. 1).

**Fig. 1.**
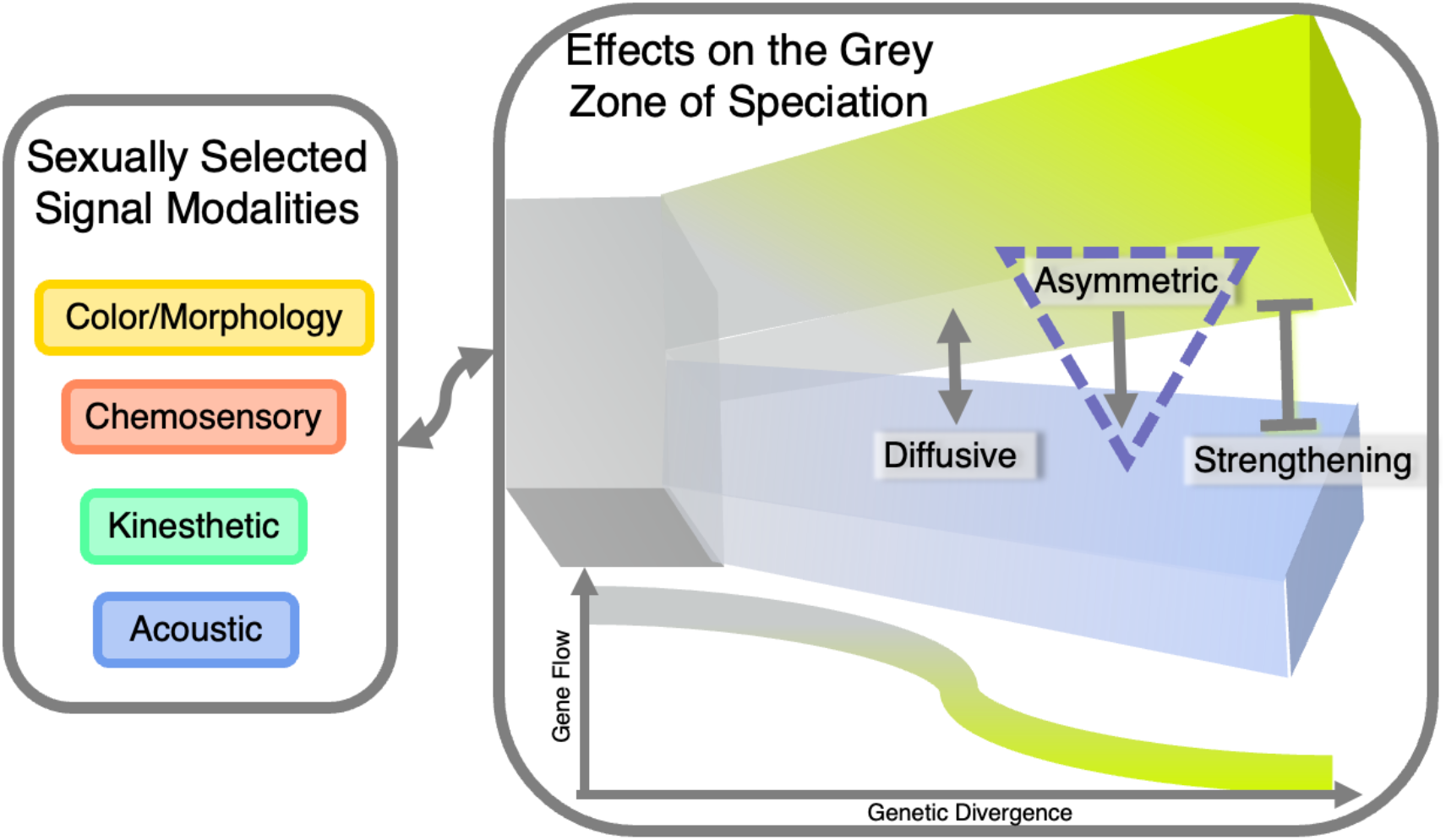
Mating signal modalities and their effects on species boundaries. Conceptual illustration showing the four mating signal modalities. Mating signals may exhibit three alternative effects of mating signals on species boundaries: diffusive, asymmetric, and strengthening.

The role of mating signals in the formation and maintenance of species boundaries has been paradoxical. Divergent mating signals often contribute to assortative mating (Jiang et al. 2013) and thus have been considered crucial for speciation (Kondrashov and Shpak 1998; Vines and Schluter 2006; Servedio 2016). However, a recent simulation study showed that assortative mating alone can be ineffective in maintaining nascent species boundaries in the face of gene flow (Irwin 2020). Even if mating signal divergence may not often be sufficient to initiate speciation, it can contribute to the speciation completion as reinforcement drives mating signal character displacement at sympatry (Noor 1999; Li et al. 2023). However, besides mediating the extent of gene flow, less is understood about how mating signals mediate the directionality of gene flow.

These mating signals can be energetically demanding (Höglund et al. 1998) to acquire and maintain on top of the energy expenditure for development (Spencer and MacDougall-Shackleton 2011), survival (Walsberg 1983; Maurer 1996), and other reproductive efforts (Welcker et al. 2015). For instance, displaying colorful plumage requires the acquisition or synthesis of pigments (Hill 1996; Jawor and Breitwisch 2003; Hill and McGraw 2006; Weaver et al. 2018). Courtship dances are energetically engaging (Barske et al. 2011; Fuxjager et al. 2022), while mating acoustics contain varying amounts of energy in sound waves (Ryan 1988; Searcy 1992; Tomaszycki and Adkins-Regan 2005).

Mating signal energetics may render diverse effects on species boundaries (Eddo et al. 2025) (Fig. 1). The relative energy expenditure of heterospecific mating signals can influence heterospecific pairing rates, as seen in multiple taxa. For example, female White-collared Manakins (*Manacus candei*) prefer the more pigmented yellow collar of Golden-collared Manakins (*M. vitellinus*) (Long et al. 2024). The more energetic songs of Collared Flycatchers (*Ficedula albicollis*) are preferred by Pied Flycatcher (*F. hypoleuca*) females (Haavie et al. 2004; Qvarnström et al. 2006). Male Peruvian Sheartails (*Thaumastura cora*) have a more complex and energy dependent courtship and mating song that has resulted in hybridization between a male sheartail and a female Chilean Woodstar (*Eulidia yarrellii*) (Clark et al. 2013). Relative energetic expenditure in mating signals can be perceived as an extended honest signal facilitating heterospecific pairing and mediating gene flow along the signal energy gradient.

Here, we conducted a data synthesis on mating signals within hybridizing avian species pairs, to answer the following questions: (1) Are mating signal modalities associated with their effects on species boundaries (Fig. 1); (2) Do mating signal energetics predict asymmetrical gene flow across species boundaries; (3) Do the species pairs with greater genetic distance have mating signals that strengthen species boundaries? Highly energetic signals serve as honest indicators of fitness (Galván et al. 2015) and may also exploit a sensory bias for complexity and novelty (Enquist and Arak 1993), stimulating mating across species boundaries. Therefore, we predict that signals with greater energetic expenditure are favored across species boundaries, leading to asymmetric gene flow along the mating signal energy gradient.

## Methods

### Hybridizing avian species pairs

To understand sexual selection energetics in the grey zone of avian speciation, we conducted a literature search. We generated metadata using the following hierarchical criteria: (1) a pair of avian species that are hybridizing; (2) within each pair, there is at least a well-documented mating signal (trait that leads to behavioral change of the receiver(s) that result in increased mate attraction and/or mate competition of the signaler) (Eddo et al. 2025); (3) the mating signal has a predicted or documented effect on hybridization.

### Mating Signal Modality

We categorized the mating signals into (1) acoustic, (2) color and/or morphology (visually received), (3) kinesthetic, (4) chemosensory (Fig. 1). A kinesthetic signal is a behavior or locomotion that facilitates gaining mates (e.g. courtship dance, bower construction, etc.) (Johnsgard 1965; Doucet 2003). Chemosensory signals are chemicals typically received through olfaction that can impact mating opportunities (Smadja and Butlin 2009). The species pairs can have multiple traits; for example, the Domestic Pigeon (*Columba livia*) and Barbary Dove (*Streptopelia risoria*) have variations in vocalizations while courting, as well as plumage color differences that are selected for (Burley 1981; Burns-Cusato and Cusato 2022).

### Mating signal effect on species boundaries

The mating signals were evaluated for their roles in the hybridizing species boundaries: (1) diffusive; (2) asymmetric; (3) strengthening (Fig. 1). A mating signal with a diffusive effect enhances heterospecific pairing equally and thus promotes hybridization. A signal with an asymmetrical effect on species boundaries favors mates from one species over another, facilitating unidirectional gene flow. A signal with a strengthening effect prevents or suppresses hybridization and reduces gene flow across the species boundaries. The signals with an asymmetrical effect on species boundaries were further categorized to best describe the gene flow. A designation of “1” was used to describe asymmetric gene flow from species 1 into species 2. A designation of “2” was used to describe asymmetric gene flow from species 2 into species 1 (Table S1).

If there was direct evidence of how this signal affected the boundary with regards to both species in a behavioral study, we listed it as *primary* evidence (Table S1). If there was evidence of both species separately, we inferred the heterospecific effect and regarded this effect as *secondary* evidence (Table S1). For cases with the understanding of the signaling function from one species and the variation of the signal expressions between species, we extrapolated the signaling effect and regarded it as *tertiary* evidence (Table S1).

### Sexual selection energetics & effects on gene flow

To assess the role of sexual selection energy in mediating gene flow across hybridizing species boundaries, we further scored the relative energetic expenditure of the mating signals that are associated with asymmetrical gene flow. For signals related to overall body mass or size, larger average body mass were considered more energetically costly (Mcnab, 1983; Taylor et al., 1982). For carotenoid-based plumage signals, which are acquired from exogenous dietary sources, red carotenoids were considered more energetic than yellow carotenoids due to the relative abundance (Hill 1996) and the enzymatic steps required to convert resources into red pigments over yellow pigments (Weaver et al. 2018). For melanin-based plumage signals, the presence of a higher concentration of melanin patches was considered of greater energy (Jawor and Breitwisch 2003; Hill and McGraw 2006). For acoustic signals, songs that had higher trill rates, higher amplitudes, and/or higher syllable diversity were considered more energetic (Christensen et al. 2006; Ritschard et al. 2010; Darolová et al. 2012; Sierro et al. 2023). Finally, for kinesthetic signals, such as courtship dances, nuptial gifts, and bower architectures, signals that required more locomotive efforts (Maurer 1996) and/or higher metabolic needs to generate these movements (Barske et al. 2011, 2014) were considered more energetically costly.

To examine whether relative energy expenditure predicts their mediation of gene flow in each hybridizing species pair, we looked further into the directionality of gene flow. For the mating signals that mediate asymmetric gene flow, we further associated the directionality of gene flow with the relative energy expenditure of the signals. A signal was assigned a score of 1 if the gene flow follows the signal energy gradient from high to low. A score of −1 was assigned to the signals with the opposite association. A signal was assigned a score of 0 if both signals required a similar level of energy. A score of NA was given if there was a lack of concrete evidence to support either side of the claim (Table S1).

### Mitochondrial genetic distance

To estimate the mitochondrial divergence between hybridizing species, we computed mtDNA distance. For each species, we downloaded sequences for three commonly sequenced mitochondrial genes—Cytochrome b (*Cytb*), NADH dehydrogenase subunit 2 (*ND2*), and Cytochrome oxidase subunit 1 (*COI*)—from NCBI GenBank for each hybridizing species in our filtered dataset using the *entrez_search* function (Winter 2017) from the *rentrez* package in R. We restricted searches to the “nucleotide” database, and to exclude non-target sequences, we included the species name followed by the “[organism]” delimiter.

We retrieved species’ sequences if at least one was available for the target gene. For species with greater than 10 sequences, we randomly selected 10 to ensure even representation of within-species diversity. We aligned sequences for each gene using MUSCLE (*msa* function; Bodenhofer et al., 2015) in R (version 4.4.2; R Core Team, 2024), with *gapOpening = 20* and *gapExtension = 10*. We then calculated genetic distances between hybridizing species using the *dist*.*dna* function with model *TN93* from the *ape* package (Paradis and Schliep 2019) and recorded pairwise mean, minimum, and maximum values for each gene. We also calculated, mean, minimum, and maximum within-species genetic distances, both per gene and as an average across genes, for comparison with pairwise divergence when at least two sequences were available. Finally, we computed the mean between and within species distance as the weighted average among estimates of all three genes. The estimates calculated from genes with greater sample sizes were weighed more.

### Phylogenetic tests

To understand the evolutionary relationship among signal modalities, their effects on species boundaries, and their association with genetic distance and the direction of gene flow between the parental species, we tested the following three associations with a phylogenetic ANOVA using the phyloANOVA function in Phytools (Revell 2024) in R. The three tests are (1) effects on species boundaries ∼ signal modality; (2) mtDNA distance ∼ signal modality; (3) mtDNA distance ∼ effects on species boundaries. For species pairs with more than one signal, we randomly selected one of the signals and combined them with the rest of the species pairs in each phylogenetic ANOVA. To understand whether the relative signal energetics are associated with the direction of asymmetric gene flow, we conducted a phylogenetic chi-square test using an equal rate model with the fitDiscrete function in R. To encompass variations of signal permutations across species pairs, we randomly conducted 1000 permutations and recorded the effect size each round.

### Avian Phylogeny

We inferred the phylogenetic relationship among all the species pairs (Fig. 2A) in our metadata from the literature. The family-level backbone of the phylogeny was derived from Stiller et al. (2024). We then refined the species-level splitting patterns with taxon-specific literature (Ellsworth et al. 1994; Johnson et al. 2001; Ksepka et al. 2006; Lovette et al. 2010; Zuccon et al. 2012; Johansson et al. 2013; Arbabi et al. 2014; Klicka et al. 2014; Smith and Clarke 2015; Dufort 2016; Sun et al. 2017; Perktaş et al. 2020; OneZoom Core Team 2021; Päckert et al. 2021).

**Fig. 2.**
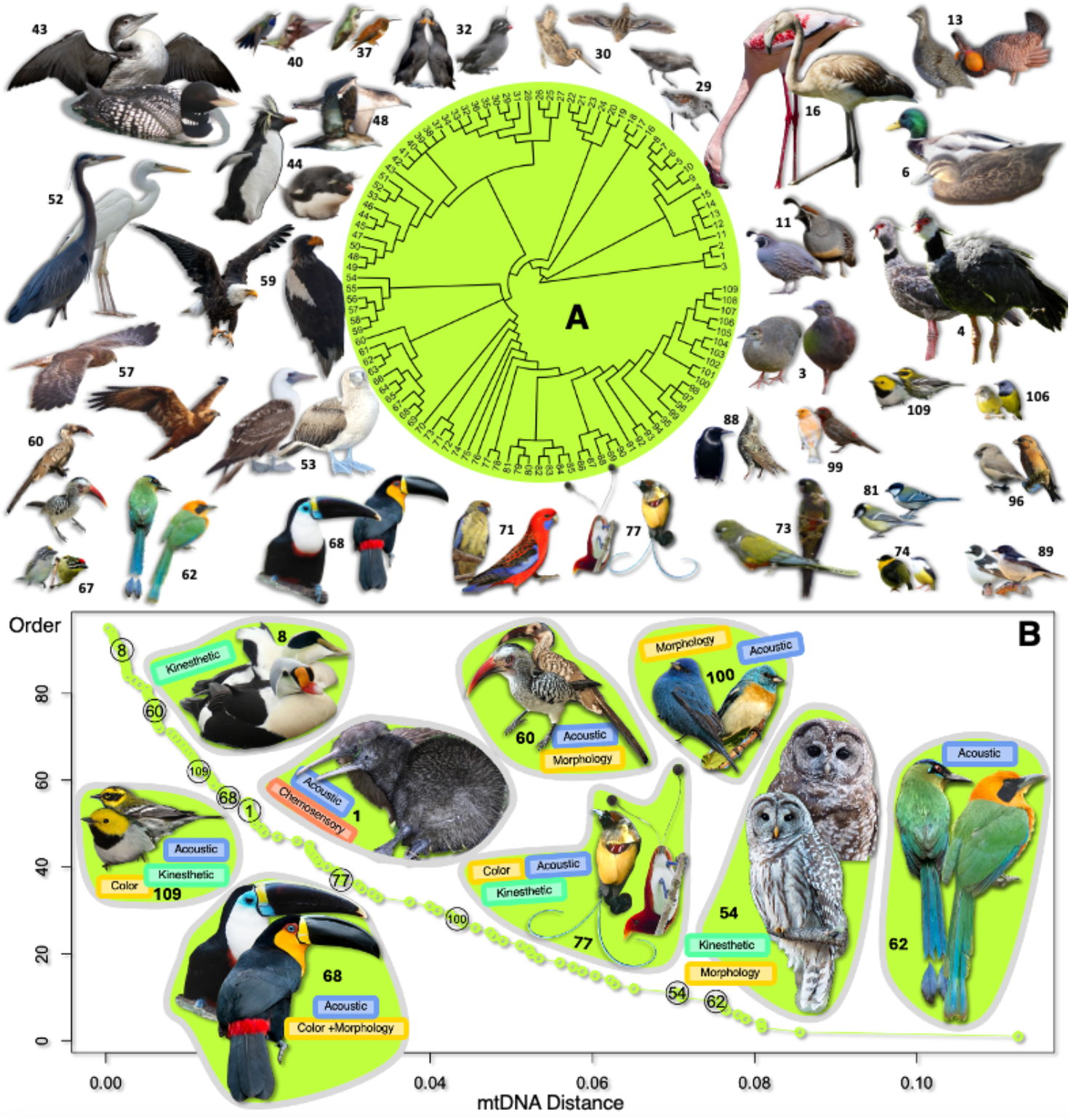
Examples of hybridizing species pairs along the speciation continuum. **A**, The 109 pairs of hybridizing species across the avian phylogeny included in this study. Details are summarized in Table S1. **B**, The genetic distance of mtDNA between hybridizing species pairs. iNaturalist photo attributions are listed in Table S2.

## Results

The complete analysis included 109 hybridizing species pairs, each containing one or more divergent mating signals (Fig. 2). The species included ranged across 26 different avian orders for a fair representation of all avian taxa. There were 51 occurrences of acoustic traits, 59 of color and/or morphology, 41 of kinesthetic traits, and 11 chemosensory cases (Fig. 3A). Among the divergent mating signals, 81 have strengthening effects, 44 are diffusive, and 38 are asymmetric (Table S1, Fig. 1).

**Fig. 3.**
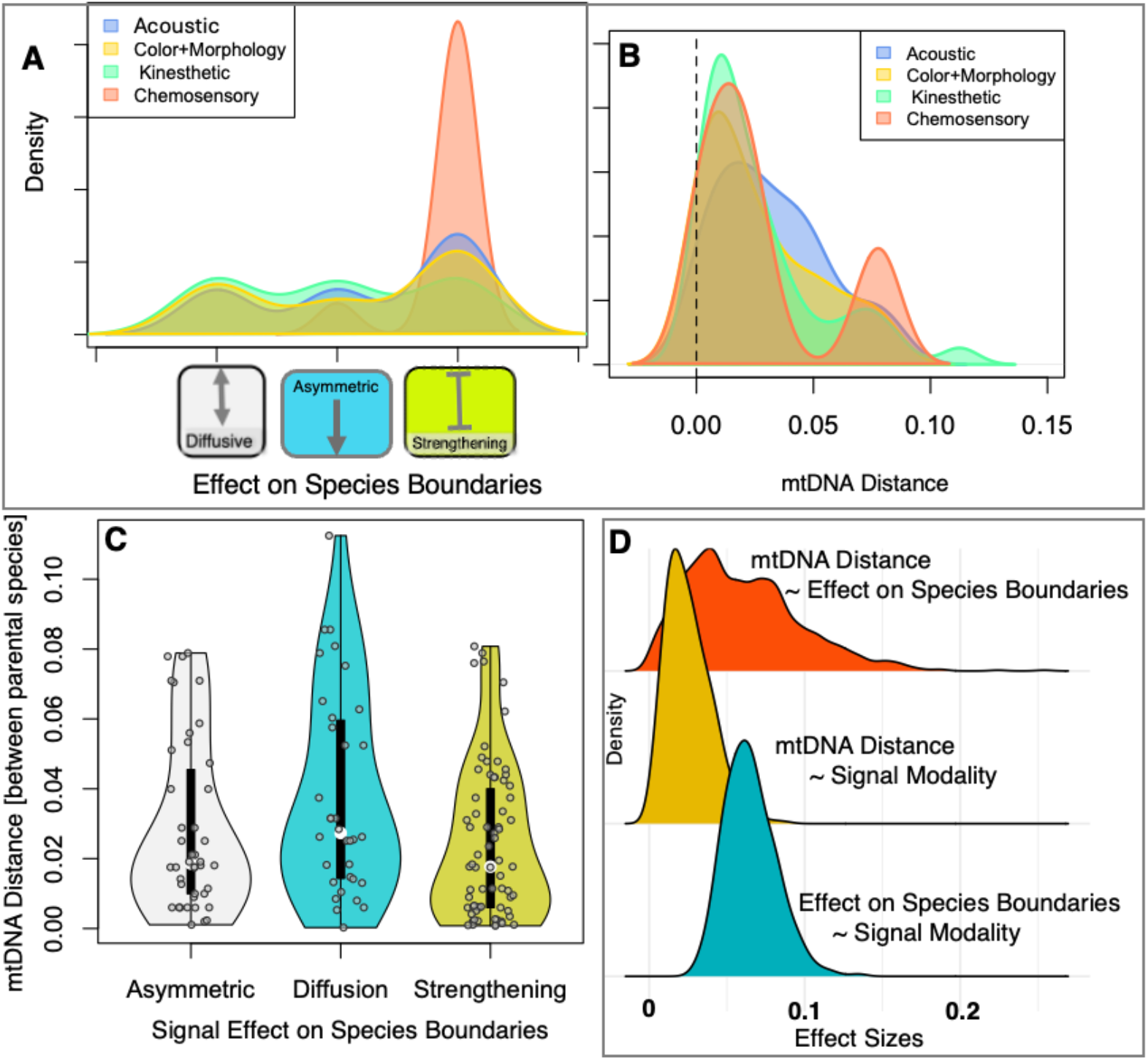
The association of signal modality, signal effect on species boundaries, and mtDNA distance between parental species. **A**, Chemosensory signals tend to strengthen species boundaries. **B**, Association between signal modality and the mitochondrial genetic distance between species. **C**, Species pairs with diffusive mating signals tend to exhibit greater mtDNA distance. **D**, The effect sizes of the phylogenetic associations (**A**-**C**) are significantly greater than zero.

We observed a significant association between mating signal modality and their effects on species boundaries (Phylogenetic ANOVA effect size = 0.065, 95%CI: 0.038 - 0.101). While the visual, acoustic, and kinesthetic signals are evenly distributed among diffusive, asymmetric, and strengthening effect categories, the chemosensory signals disproportionately exhibited strengthening effects (Fig. 3 AD). Mating signal modality also showed a significant but weak association with the mtDNA distance between the parental species (Phylogenetic ANOVA effect size = 0.026, 95% CI: 0.005 - 0.061) (Fig. 3 BD). The species pairs with diffusive instead of strengthening signals harbor greater mtDNA distance (Phylogenetic ANOVA effect size = 0.017, 95% CI: 0.0005, 0.0556) (Fig. 3 CD).

Among the species pairs demonstrating asymmetric gene flow, the mating signal energy gradient significantly predicted the directionality of that gene flow. In particular, the gene flow direction predominantly goes from species with high-energy mating signals to species with low-energy mating signals (Phylogenetic *χ*^2^= 20.58, *p* < 10^−5^; Fig. 4).

**Fig. 4.**
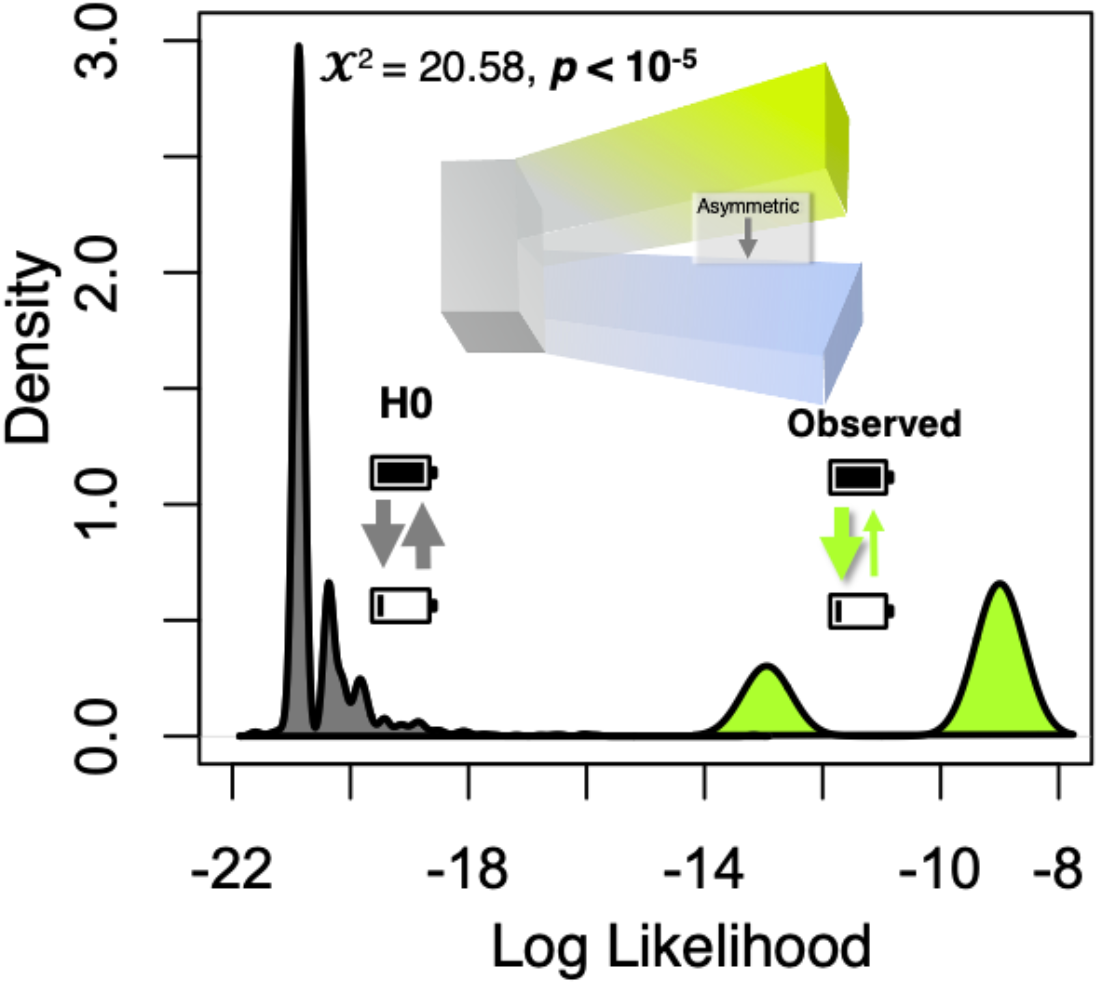
Asymmetric gene flow follows mating signal energy gradient. The observed phylogenetic association between heterospecific mating asymmetry and relative mating signal energetics is greater than random association (*p* < 10^−5^). The grey density represents the log likelihood distribution of the null model. The bright yellow distribution represents the log likelihood distribution of observed.

## Discussion

### Mating signal & speciation

Our results showed that species pairs with greater mitochondrial divergence tend to exhibit diffusive rather than strengthening mating signals (Fig. 3). The species pairs with greater mtDNA distance and diffusive mating signals often possessed one or few hybrid observations, e.g. Hairy (*Dryobates villosus*) and Ladder-backed (*D. scalaris*) Woodpeckers (Miller 1955); Rockhopper (*Eudyptes chrysocome*) and Macaroni (*E. chrysolophus*) Penguins (Woehler and Gilbert 1990); and Rufous (*Baryphthengus martii*) and Amazonian (*Momotus momota*) Motmots (Marcondes et al. 2013). Many such species pairs experienced periods of allopatry, which is recognized as the most common geographic mode of speciation in birds (Price 2008). Geographically isolated species preclude the evolution of divergent mating signals when hybridization is rare. In contrast, sympatric species typically have greater species-specific mating discrimination, reinforcing sexual isolation rather than physical (Noor 1999). This pattern is evident in *Ficedula* flycatchers, where sympatric populations within the hybrid zone of Pied (*F. hypoleuca*) and Collared Flycatchers (*F. albicollis*) displayed greater plumage color divergence than allopatric populations of this species pair (Sætre et al. 1997).

While premating isolation, such as species-specific mating discrimination, is important in maintaining species boundaries, it is often singly insufficient for speciation by limiting gene flow (Irwin 2020). However, assortative mating is often accompanied by postzygotic isolation such as reduced hybrid fitness (McQuillan et al. 2018; Alario et al. 2023; Blain et al. 2024), which reinforces the effects of premating isolation to drive speciation. This process is more pronounced in sympatric species, where both premating and postmating isolation are stronger due to higher hybridization risks (Ball and Jameson 1966). In contrast, when hybridization is rare, there is insufficient variation for reinforcement to diverge premating signals to prevent hybridization (Pyron and Burbrink 2010). This process likely leads to our observation (Fig. 4 CD), with rare hybridization events stemming from extended geographical isolation with less reliance on divergent mating signals.

### Gene flow along mating signal energy gradient

We observed significant directional gene flow following the mating signal energy gradient (Fig. 4). This is likely due to higher energy expenditures accurately showing greater fitness in many species pairs, described in the literature as signal honesty (Grafen 1990; Fromhage and Henshaw 2022). These signals often require a higher cost or energy input, such as higher rates of aggression between courting males (Augustine and Trauba 2015) or longer and high-frequency songs (Buchanan and Catchpole 1997) leading to a higher rate of mating success.

### Acoustic signal energetics

Avian acoustic signals are quantified in terms of frequency, amplitude, syllable diversity, trill rate, and duration—all of which depend on energetic composition (Gil and Gahr 2002; Ballentine et al. 2004; Nowicki and Searcy 2005). These vocal traits vary in their costs and functions, influencing mating dynamics in species across different contexts (Catchpole 1980; Eens et al. 1991). Frequency and amplitude require greater muscle effort and lung capacity, with the relatively higher-value sounds requiring more energy (Oberweger and Goller 2001). Frequency is primarily controlled by syringeal muscles, where activity increases exponentially with song frequency, while higher amplitude demands greater oxygen consumption and metabolic effort (Oberweger and Goller 2001). Syllable diversity, being indicative of cognitive ability and genetic quality, requires more vocal effort and coordination, thus requiring more energy (Barnett and Briskie 2007). Fast and sustained trills impose an energetic burden due to respiratory effort and air pressure depletion (Sierro et al. 2023), indicating that their combination with other traits can further increase energy demands. Similarly, longer song duration requires sustained energetic input, requiring at least a moderate increase in metabolic rate (Ward et al. 2003). These traits often exist in hierarchical trade-offs, balancing not only with each other but with other fitness signals such as body condition or plumage quality as well (Ballentine et al. 2004; Cardoso et al. 2007).

Because these traits serve as honest indicators of fitness, they also influence species recognition and mate selection, sometimes contributing to hybridization when one species has more desirable song characteristics (Eens et al. 1991; Buchanan and Catchpole 1997; Ballentine et al. 2004). In some cases, females select mates based on performance rather than species identity. This is also observed in a well-studied hybrid zone between Pied (*Ficedula hypoleuca*) and Collared (*F. albicollis*) Flycatchers, where the song of the Collared Flycatcher was preferable to that of the Pied (Qvarnström et al. 2006). The Collared Flycatcher’s songs are more energetically demanding with longer duration and higher frequency (Haavie et al. 2004) leading to directional gene flow.

Similarly, the energetic calls of Damaraland Red-billed Hornbills (*Tockus damarensis*) with higher frequencies and syllable diversity introgress asymmetrically into the range of Northern Red-billed Hornbills (*T. erythrorhynchus*) (Delport et al. 2004).

### Visual signal energetics

There are multiple cases where an asymmetric gene flow follows a body size gradient. For example, female Clapper Rails (*Rallus crepitans*) preferentially mated with males of the much larger King Rail (*R. elegans*) (Coster et al. 2018). This pattern was seen in taxa with reversed sexual size dimorphism. In the Lesser (*Clanga pomarina*) and Greater Spotted Eagles (*C. clanga*), males of the former preferentially mated with females of the latter due to their larger body size (Väli et al. 2010). This body size gradient is also observed in Jacanas, a sex-role reversed system, in which the larger body size of the Northern Jacana (*Jacana spinosa*) introgresses into the smaller Wattled Jacana (*J. jacana*) (Lipshutz 2017). The body size differential between hybridizing species likely predicts asymmetric gene flow, as a larger body size is perceived as an honest signal (Taylor et al. 1982; Mcnab 1983).

Color energetics is also associated with the direction of gene flow between species. Female Peruvian (*Sula variegata*) Boobies pair with the pigmented Blue-Footed Boobies (*S. nebouxii*), but reciprocal pairing is less likely (Taylor et al. 2010). This pattern aligns with a bias for red pigmentation that has been observed across several hybridizing species. Between Red-fronted (*Pogoniulus pusillus*) and Yellow-fronted (*P. chrysoconus*) Tinkerbirds, females of both species have a mating preference for male red-fronted tinkerbirds, likely due to their red plumage coloration (Nwankwo et al. 2019). Additionally, female Orange-backed Fairywrens (*Malurus melanocephalus melanocephalus*) often have extra-pair copulations with male Red-backed Fairywrens (*M. m. cruentatus*) (Baldassarre and Webster 2013). Red coloration has been stated to be more energetically rich than yellow hues, as there are more conversion steps within the carotenoid process (Weaver et al. 2018). The relative abundance of red carotenoids may also be limited, requiring significant foraging effort to obtain (Hill 1996). Overall, pigmentation that requires energy to forage, metabolize, implement, and maintain is likely to act as an extended honest signal during heterospecific mating (Galván et al. 2015).

### Kinesthetic signal energetics

Asymmetric gene flow follows the same energy gradient seen above in the kinesthetic signaling modality. In the mating interactions of Golden-collared (*Manacus vitellinus*) and White-collared Manakins (*M. candei*), there is courtship dominance of the male former over the latter (McDonald et al. 2001; Stein and Uy 2006; Barske et al. 2014, 2023; Bennett et al. 2021; Long et al. 2024). Golden-collared Manakins display their preferential yellow collar during courtship displays in lekking arenas, giving them an advantage over White-collared males (Barske et al. 2014, 2023). Golden-collared males are more aggressive than those with a white collar, highlighted by their likelihood to attack mounts, leading to further Golden-collared male dominance (McDonald et al. 2001). On top of aggressive behaviors, Golden-collared Manakins display a greater kinesthetic effort during courtship lekking (Barske et al. 2014), which may lead to preferential mating from females (Barske et al. 2011). In heterospecific lekking arenas, females of both species preferentially mate with male Golden-collared Manakins, often rejecting White-collared males (Stein and Uy 2006). Both the overall higher energy expenditure and aggressive nature of Golden-collared Manakins likely contribute to their mating success, as females prefer males who express quality traits through their kinesthetic mating signals (Barske et al. 2011; Galván et al. 2015).

### Signal modalities and speciation effects

Mating signal modalities are significantly associated with their effects on the species boundaries (Fig. 3 AD). This association is mainly driven by the disproportionate rate at which chemosensory signals had a strengthening effect. Kinesthetic, visual, and acoustic signals had a relatively even distribution among the diffusive, asymmetric, and strengthening effect categories (Fig. 3 A).

### Chemosensory signals

While chemosensory signals remain relatively understudied in birds, our findings suggest they could be a key factor in maintaining species boundaries. Several studies indicate that individuals prefer conspecific chemosensory cues, suggesting a significant role for this modality in strengthening species boundaries (Whittaker et al. 2011; Krause et al. 2023). In a key study, it was found that in the hybridizing complex between the Black-capped (*Poecile atricapillus*) and Carolina (*Poecile carolinensis*) Chickadees, both males and females preferred conspecific scents, leading to heterospecific premating isolation (Van Huynh and Rice 2019). The two species varied in their preen oil compositions, with the Carolina Chickadee having a higher ratio of ester to non-ester compounds (Van Huynh and Rice 2019). Although the specific functions of these compounds in chickadees remain unclear, esters are known to aid in feather cleanliness, provide waterproofing, offer potential protection against bacteria and parasites, and enhance mate attractiveness during the breeding season (Haribal et al. 2005). The difference in preen oil composition possibly stems from the divergent environmental conditions of the two chickadee species that breed in distinct latitudes (Van Huynh and Rice 2025). However, further research is needed to evaluate the potential connection between chemosensory compound roles and local adaptation. Given that hybrids in this complex exhibit reduced cognitive function and hatching success (Bronson et al. 2005; McQuillan et al. 2018; Van Huynh and Rice 2019), these chemical differences may aid species boundary reinforcement by signaling hybrid incompatibilities.

Alternatively, an opposite effect of chemosensory signaling on species boundaries may occur when it is linked to Major Histocompatibility Complexes (MHCs). Preen oil’s biochemical composition has been linked to fitness through MHCs. The genetic diversity in MHCs is often positively correlated with immune system function (Spurgin and Richardson 2010). Furthermore, a recent comparative study of animal hybrid zones revealed excessive introgression of MHC genes relative to genome background (Gaczorek et al. 2024). Chemosensory signals have been shown to carry information on MHCs (Leclaire et al. 2017; Jennings et al. 2022), guiding disassortative mating in MHCs.

Although research on the energetic aspects of chemosensory signaling in birds is limited, this phenomenon has been explored in other species, such as mice. It has been found that the creation of urinary peptides, known to be used in chemosensory signaling for both mate attraction and territoriality, involves complex biochemical pathways that require great amounts of ATP (Brennan and Kendrick 2006). It has also been discovered that olfactory sensory neurons used to detect and process odors in various species also rely on ATP to mobilize CA^2+^ for this process (Fluegge et al. 2012). The core chemosensory energetic pathways may be conserved in different endotherms and influence interspecific/intraspecific mating interactions. Collectively, chemosensory signals may be linked to different fitness components, eliciting a complex mate choice dynamic at species boundaries.

## Future directions

Mating signals are often hierarchical, with multiple traits differing in their relative importance to mate choice and reproductive success. This is evident in plumage coloration among Psittaciformes, where pigmentation and structural mechanisms interact to produce distinct hues. Parrot feather coloration is generated through psittacofulvins, which synthesize yellow pigments via aldehyde dehydrogenase ALDH3A2 (Arbore et al. 2024), and through structural modifications that reflect light to produce blue hues (Prum et al. 1998). The combination of these mechanisms yields green plumage (Arbore et al. 2024), illustrating the interplay of multiple developmental pathways. Similarly, in Bowerbirds, mating signaling extends beyond morphological traits to elaborate structural displays. Males invest significant energy in foraging for materials and constructing bowers, a process that requires both sustained effort and resource acquisition. This multi-step investment exemplifies how complex signaling traits can be shaped by energetic constraints (Doucet 2003). Quantifying the relative energetic costs of each component—bower construction, maintenance, and display— remains a challenge. Additionally, sexually selected traits can have secondary functions, such as melanin pigmentation contributing to parasite resistance (Gasparini et al. 2009; Jacquin et al. 2011) or the association between high leukocyte counts and red psittacofulvin concentrations in parrots (Edwards 2012). Future research integrating these physiological and behavioral dimensions could yield a more comprehensive framework for understanding the energetic trade-offs in mating signaling.

## Conclusion

Our results are consistent with the rising understanding that mating signals which strengthen species boundaries are not necessarily concordant with the greater genetic divergence that had grown predominantly in allopatry. Among the species pairs with great genetic divergence but limited hybridization, the efficacy of reinforcement is limited to selecting mating signals that strengthen species boundaries. Although mating signals might be insufficient in initiating speciation alone, they have a significant effect on the directionality of gene flow across nascent species boundaries. Here, we highlight the importance, yet limited understanding, of mating signal energetics, which may play a role in asymmetric gene flow across species boundaries. Understanding the energetics behind the plethora of colorful, melodious, aromatic, and locomotive signals of the arial tetrapods shed light on the origin and maintenance of species boundaries.

## Acknowledgment

We are thankful to Ummat Somjee, Michael Ryan, Justin Havird, Dahong Chen, and Silu Wang’s Lab for helpful discussions. Funding is provided by the SUNY Research Foundation startup grant awarded to SW.

